# Mosquito Transmission and Human Hepatocyte Infections With *Plasmodium ovale curtisi* and *P. ovale wallikeri*

**DOI:** 10.1101/435768

**Authors:** Mojca Kristan, Samuel G. Thorburn, Julius Hafalla, Colin J. Sutherland, Mary C. Oguike

## Abstract

Human ovale malaria is caused by the two closely related species *Plasmodium ovale curtisi* and *P. ovale wallikeri*. Both species are known to relapse from quiescent hepatic forms months or years after the primary infection occurred. Although some studies have succeeded in establishing mosquito transmission for ovale malaria, none have specifically described transmission and human hepatocyte infection of both sibling species. Here we describe a simplified protocol for successful transmission of both *P. ovale curtisi* and *P. ovale wallikeri* to *Anopheles coluzzii* mosquitoes, and streamlined monitoring of infection using sensitive parasite DNA detection, by loop-activated amplification, in blood-fed mosquitoes. In one experimental infection with *P. ovale curtisi* and one with *P. ovale wallikeri*, viable sporozoites were isolated from mosquito salivary glands, and used to successfully infect cultured human hepatocytes. This protocol provides a method for the utilisation of pre-treatment clinical blood samples from ovale malaria patients, collected in EDTA, for mosquito infection studies and generation of the hepatic life cycle stages of *P. ovale curtisi* and *P. ovale wallikeri*. We also demonstrate the utility of LAMP as a rapid and sensitive alternative to dissection for estimating the prevalence of infection in *Anopheles* mosquitoes fed with *Plasmodium*-infected blood.

## INTRODUCTION

*Plasmodium ovale* is a parasite causing human malaria, with a wide distribution across tropical regions, especially sub-Saharan Africa and some islands of the western Pacific [1, 2]. It was described as a species in 1922 [1], while the two forms *P.ovale curtisi* and *P.ovale wallikeri*, first distinguished in 2010, are now regarded as separate species [2, 3]. All human *Plasmodium* parasite infections are initiated by a bite from an infected mosquito, injecting sporozoites that invade hepatocytes and form an obligatory liver stage, which develops into mature intra-hepatocytic schizonts. In *P. vivax*, *P.o. curtisi* and *P.o. wallikeri*, a proportion of sporozoites undergo developmental arrest to form latent liver-stage forms called hypnozoites, which resume hepatic development, due to unknown triggers, and cause relapses months or years after the initial infection [4–9]. The formation of hypnozoites is of public health importance, as these forms are long-lasting reservoirs of infection.

Artemisinin combination therapy (ACT) and other schizonticidal antimalarial drugs can clear blood-stage *P. ovale* spp. infections, but the 8-aminoquinoline primaquine is the only drug currently in use that can kill hypnozoites [5]. Malaria elimination will depend on such anti-hypnozoite medicines to prevent relapses, however primaquine-based therapy has three major drawbacks [10]. Primaquine causes haemolysis in G6PD-deficient individuals, it has a short half-life, and as such must be administered for up to 14 days, and screening populations for G6PD deficiency prior to primaquine administration is usually not feasible [11]. New drugs, such as tafenoquine [12], are being developed but studies of the susceptibility of *P. ovale* spp. hypnozoites to these compounds require infectious gametocytes for generation of sporozoite-positive mosquitoes, and for subsequent hepatic-stage growth *in vitro*. The lack of a continuous culture system for *P. ovale* spp., due to a requirement for reticulocytes for parasite development [1], means that ovale gametocytes for generation of sporozoites, and thus hepatic stages, can only be obtained from human infections. These obstacles have prevented much progress in drug susceptibility studies of the hepatic stages of *P. ovale* spp. [12].

The Malaria Reference Laboratory (MRL) at LSHTM receives blood samples from UK cases of imported malaria for specialist diagnosis, and surplus diagnostic material is available for further studies [2, 7]. Our aim was to develop a protocol using the available EDTA-preserved blood samples from patients infected with *P. ovale* spp. for infective membrane feeds; producing sporozoites for use in hepatocyte invasion assays; and carrying out drug susceptibility studies of liver-stage parasites. We describe proof-of-principle transmission of *P. ovale* using up to 4 days old EDTA-preserved blood samples, subsequent penetration of sporozoites into hepatocytes and evidence of successful differentiation into exoerythrocytic forms (EEF).

## METHODS

### *Plasmodium ovale* samples and mosquito infections

Anonymised, *Plasmodium*-infected blood samples from the Public Health England Malaria Reference Laboratory were obtained for the purpose of developing a work-flow suitable for evaluating liver-stage parasite susceptibility to antimalarial drugs. Sodium EDTA-preserved blood samples from gametocytaemic patients infected with mono-infections of *P. o. curtisi* or *P. o. wallikeri* were mixed 1:1 with fresh blood from donors (blood group O; anonymised; collected under a protocol approved by the LSHTM Research Ethics Committee) and used for membrane feeding. Parasite densities in the patient blood samples were estimated *post hoc* from Giemsa-stained thin blood films, read by a single microscopist.

Female *Anopheles coluzzii* (N’gousso strain [13]), two to six days post-emergence, were used in the experiments. The mosquitoes were kept in an incubator at 27°C and 70% relative humidity throughout the experiment. Mosquitoes were given *ad libitum* access to 10% glucose/0.05% PABA (para-aminobenzoic acid) solution from 24 hours after the feed, so that unfed mosquitoes died during this first 24 hours.

Daily mosquito mortality was recorded. Mosquitoes were dissected 15-19 days after the infective feed. Prior to dissection, the insects were anaesthetised using ethyl acetate and kept on ice. Each mosquito was transferred to a small petri dish containing 70% ethanol, and then to another containing RPMI medium. Salivary glands were dissected from mosquitoes and kept in RPMI, on ice.

### Sporozoite hepatocyte invasion assay

Three different hepatocyte lines (Huh7, IHH and HepG2) were tested for optimal sporozoite invasion rates by plating out different numbers of cells (30,000; 20,000 and 10,000) in Labtek wells. *P. falciparum* sporozoites were used for testing various hepatocyte culture conditions (*P. falciparum-*infected *An. stephensi* mosquitoes were a kindly donated by colleagues at Imperial College, London). Cells were incubated for 48, 72, 96 and 144 hours to observe the growth of exoerythrocytic forms (EEF).

*P. ovale* spp. sporozoites were plated onto the human hepatocyte line Huh7, cultured in RPMI complete (RPMI [Gibco] with 10% foetal calf serum [Gibco], 2% L-glutamine [Gibco] and 1% penicillin/streptomycin [Gibco] at 37°C with 5% CO_2_). 24 hours before sporozoite invasion, cells were seeded into Labtek wells (ThermoFisher Scientific) at 30,000 cells per well. On the day of infection salivary glands were dissected from *P. ovale* spp.-infected female *An. coluzzii* mosquitoes. Glands were homogenised and sporozoites isolated by centrifugation. In each experiment, all sporozoite material was added to one well on the Labtek slide as sporozoite counts were low (below 2,000 in total). Slides were centrifuged and incubated for 2 hours at 37°C with 5% CO_2_ to permit hepatocyte invasion before being washed 3 times with RPMI complete. Slides were incubated for a further 3 to 6 days to allow for EEF development. To stain, cells were fixed with 4% paraformaldehyde (PFA) and permeabilised with 0.3% Triton X-100 (Sigma). EEF were stained with rabbit antibodies specific to *P. falciparum* heat shock protein 70 (PfHSP-70) (StressMarq) at 1:50 dilution, followed by goat anti-rabbit IgG Alexa Fluor 488 fluorescent antibodies (ThermoFisher Scientific) at 1:5000 dilution. Nuclei were stained with DAPI (ThermoFisher Scientific). Cells were coated in Vectashield (Vectashield^®^ Mounting Medium, Vector Laboratories) under a sealed coverslip. Imaging was performed on a Zeiss LSM510 confocal microscope or a Nikon Eclipse Ti-E fluorescent microscope.

### Loop-mediated isothermal amplification (LAMP) for detection of parasites in mosquitoes

DNA was extracted from fed mosquitoes by grinding the mosquitoes using plastic pestles then boiling in 0.1mL of 1×TE buffer (10mM Tris-HCl, pH 8.0, 1mM EDTA), followed by standard phenol/chloroform/isoamyl alcohol extraction [14]. Pan-*Plasmodium* LAMP (Eiken Chemical Co., Tokyo, Japan) [15] was used according to manufacturer’s instructions to amplify genus-specific mitochondrial target sequences as confirmation of infection status of all mosquitoes. Presence of parasite DNA in mosquitoes at least 5 days post-feed was considered evidence of an established *P. ovale* spp. infection.

## RESULTS

### Suitability of different hepatocyte lines

Multiple cell lines have been used in order to create conditions that promote *in vitro* hepatocyte invasion by malaria parasites, including HepG2 and Huh7 lines for human-infecting *Plasmodium* species. In preliminary experiments, we compared HepG2, Huh7 and the IHH cell line (Kings College, London) using *P. falciparum* sporozoites from NF54-infected mosquitoes. When either IHH or Huh 7 cells were incubated for 96 hours with *P. falciparum* sporozoites, EEF were observed developing in host hepatocytes. No EEF were observed with HepG2 cells. Parasites in the IHH cells looked rounded while those in the Huh7 cells appeared to be larger and better differentiated, an indication of more successful *in vitro* EEF development (data not shown). Seeding of hepatocytes at 30,000 per well was associated with optimal *P. falciparum* EEF invasion and development, hence 30,000 Huh7 cells per well were plated out for *P. ovale* spp. invasion assays.

### *Plasmodium ovale* spp. transmission experiments

Six infective feeding experiments were carried out between April 2016 and March 2017, using anonymised PCR-confirmed *P. ovale* spp.-infected patient blood (obtained from the Malaria Reference Laboratory, LSHTM) in EDTA. Details of parasite density and gametocyte carriage are presented in Table 1.

**Table 1.** Overview of the experiments and material used.

| Experiment | Parasite species | Parasite origin | Parasite density*<br>*remarks* | Gametocytes<br>present** | Day of salivary gland<br>dissection |
| --- | --- | --- | --- | --- | --- |
| 1 | *P.o. wallikeri* | Nigeria | 78 | + | 15 days |
| 2 | *P.o. wallikeri* | Nigeria | 338 | +++ | 18 days |
| 3 | *P.o. curtisi* | Nigeria | 48<br>*mostly late stages* | +++ | 19 days |
| 4 | *P.o. wallikeri* | Nigeria | 120<br>*early stages* | + | 19 days |
| 5 | *P.o. wallikeri* | Not specified | 22 | + | 16 days |
| 6 | *P.o. curtisi* | Nigeria | 28<br>*late stages only* | + | 15 days |

\* All stages *P. ovale* spp. per 100 leukocytes on Giemsa stained thin blood film
\*\* Present: +; Frequent: ++; Abundant: +++

Mosquito survival after the infectious feed was monitored. Excluding the non-fed mosquitoes, which all died in the first 24 hours, 40.4%, 34.2% and 32.9% of the fed mosquitoes survived in the first, second and third experiments, respectively (Fig. 1). Survival in different feeds did not vary significantly (Log-rank test, *p* = 0.5928).

**Figure 1.**
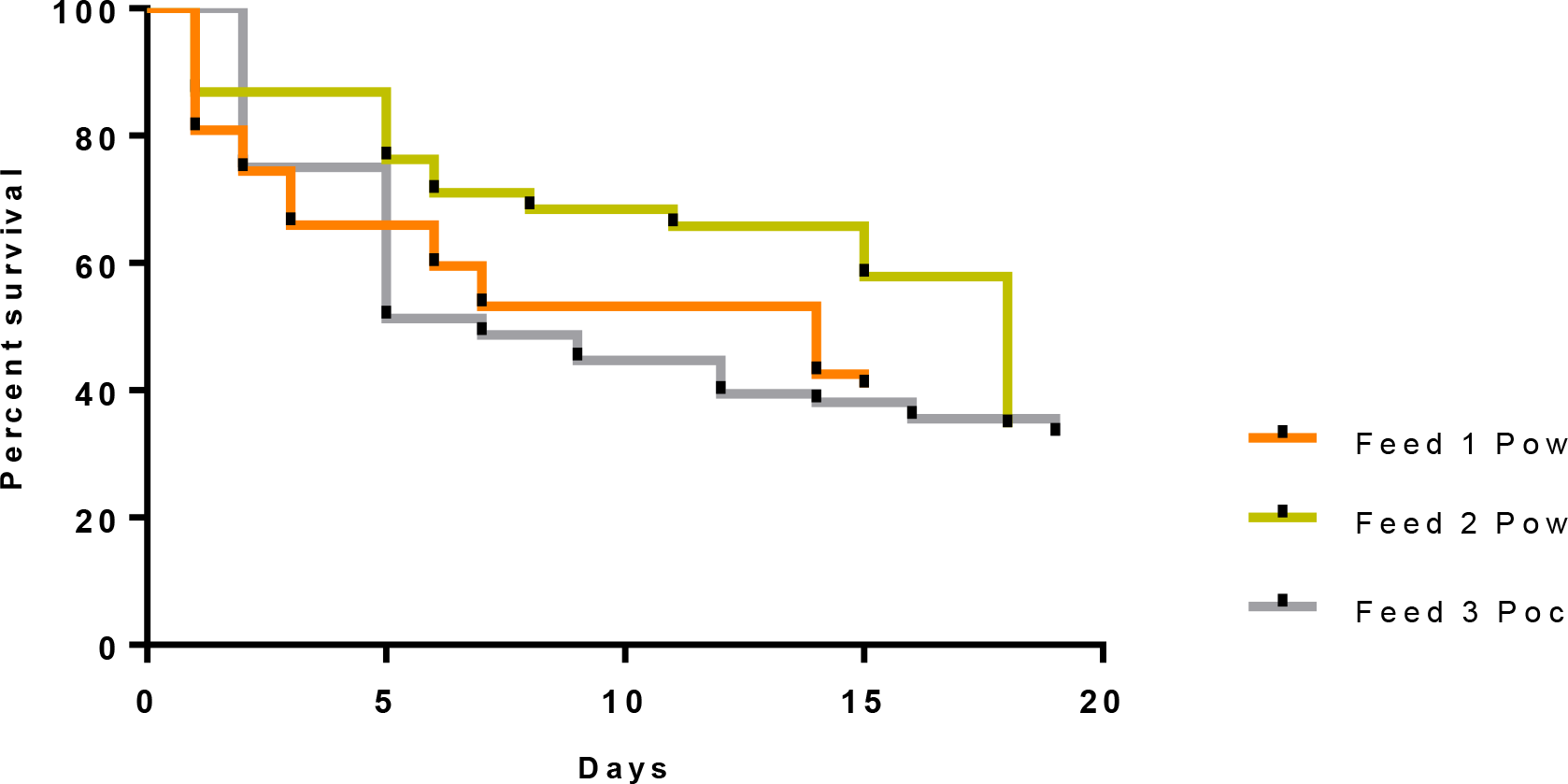
Mosquito survival following transmission of *P. ovale* spp. to *An. coluzzii* in 3 experiments. The percent survival of blood-fed mosquitoes after an infective feed is plotted over time (days). Starting numbers of mosquitoes in each experiment were 70, 60 and 100, respectively.

Parasite DNA was detected in mosquitoes from each of feeding experiments 1 to 3, but not in experiments 4 to 6 (Table 2). These unsuccessful experiments are not considered further here. A high proportion of mosquitoes in the group that died within the first two days following the feed were positive for parasite DNA by LAMP (Fig. 2). However, this is likely to reflect DNA present in the blood meal and not an established viable infection. Parasite DNA was also detected in mosquitoes at least 5 days post-feed, indicating established infection, in each of the first three feeds (Table 2; Fig. 2). Sporozoite DNA was detected in salivary gland material from experiments 1 and 3, with 10.5% (2/19) and 12% (3/25) of mosquitoes LAMP positive (Fig. 2). Sporozoites were observed by microscopy, at low density, in pooled salivary gland homogenates from feeds 1 and 3 (Fig. 3). Using haemocytometer counts, 2000 sporozoites were isolated from feed 1; however numbers were too low to accurately estimate sporozoite numbers for feed 3.

**Table 2.** Summary of successful transmission experiments.

| Expt | Species | No. mosquitoes offered a blood meal | No. mosquitoes blood-fed | % LAMP positive (n)* | % established infection (n) | Number. dissected | Number LAMP-positive | Sporozoites observed |
| --- | --- | --- | --- | --- | --- | --- | --- | --- |
| 1 | *P.o. wallikeri* | 70 | 48 | 37.5 (18) | 16.7 (8) | 19 | 2 | YES |
| 2 | *P.o. wallikeri* | 60 | 38 | 15.8 (6) | 7.9 (3) | 13 | 0 | - |
| 3 | *P.o. curtisi* | 100 | 76 | 27.6 (21) | 7.9 (6) | 25 | 3 | YES |

\*n = number of fed mosquitoes retrieved from day 5 post-feed onwards and tested by LAMP

**Figure 2.**
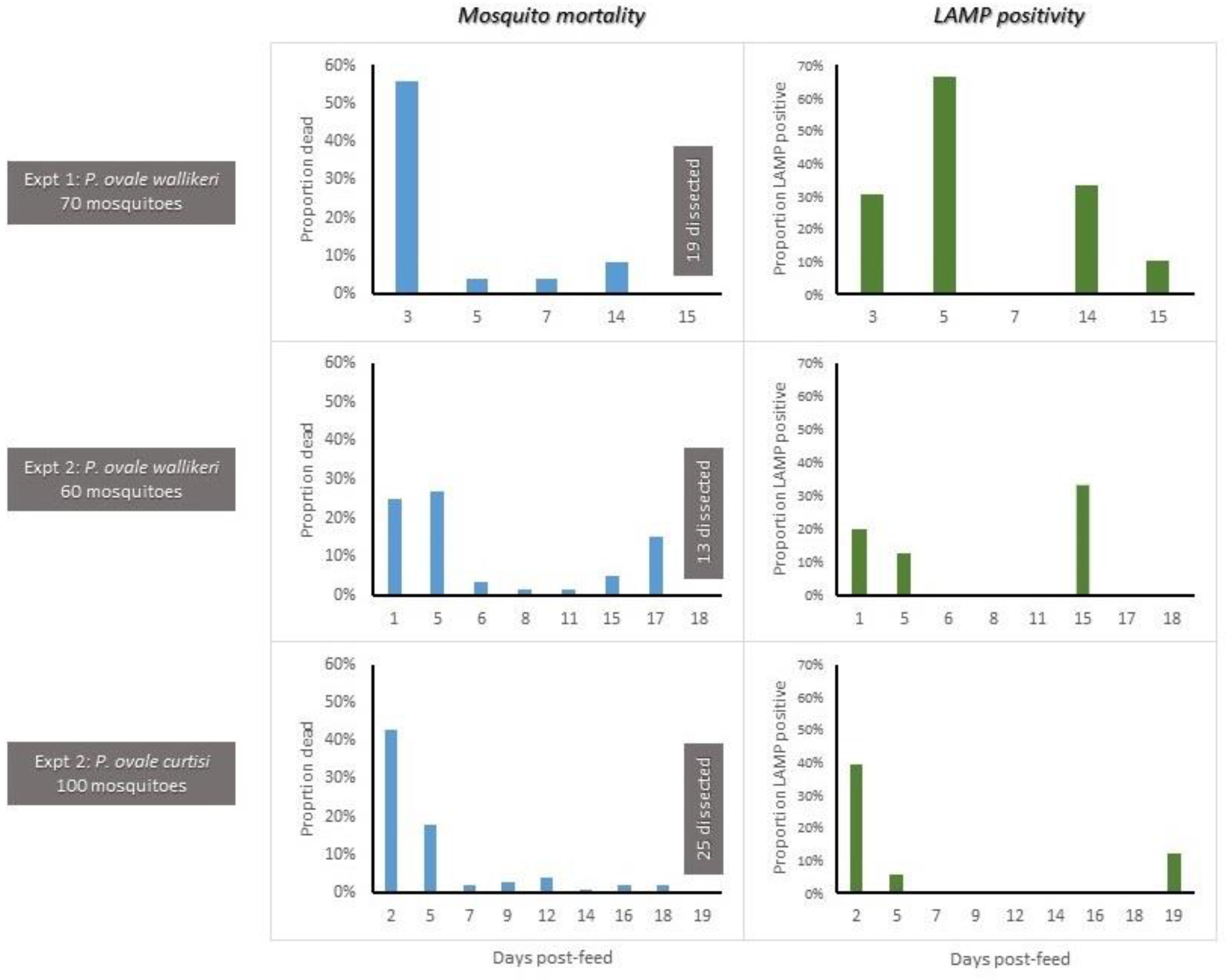
Detection of *P. ovale* spp. DNA in blood-fed *An. coluzzii*. For three experimental transmissions conducted with *P. o. wallikeri*, *P. o. wallikeri* and *P. o. curtisi*, respectively, dead mosquitoes collected on any day and those surviving to dissection on the days indicated were tested for the presence of *Plasmodium* DNA by LAMP. Sporozoites were successfully harvested on day 15 in experiment 1, and on day 19 in experiment 3.

**Figure 3.**
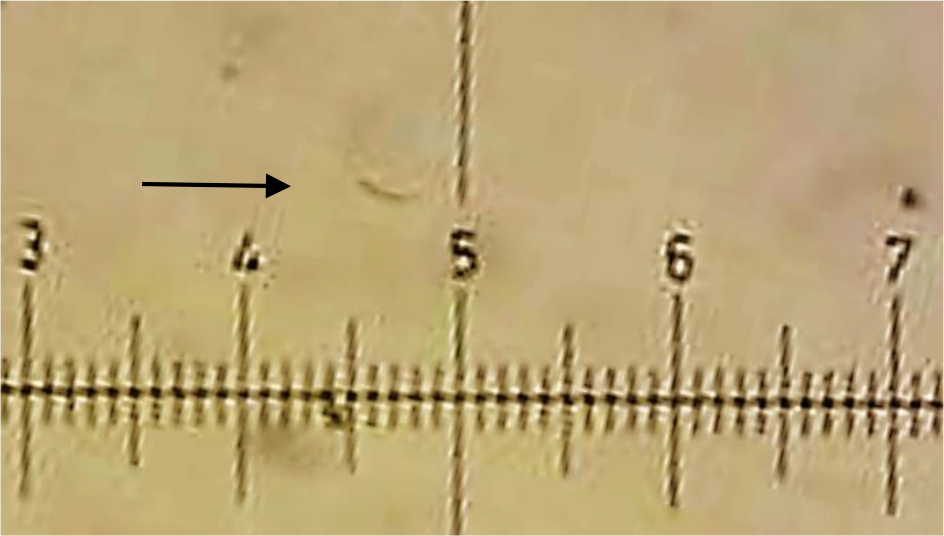
*P. ovale wallikeri* sporozoite. Bright field image of a single, motile sporozoite (*arrow*) taken at ×20 magnification after isolation from day 15 infected female *An. coluzzii* mosquito salivary glands (experiment 1).

### *Plasmodium ovale* spp. hepatocyte invasion assay

In order to investigate the capacity of the sporozoites to successfully invade hepatocytes, material isolated from all salivary glands dissected from each experiment was pooled and added into a single well seeded with Huh7 cells. After incubating 3-6 days for intrahepatic development, EEF were visualised by PfHSP-70 fluorescence in juxtaposition with DAPI-stained host-cell nuclei. Reflecting the low numbers of sporozoites, a single EEF was observed in experiment 1, and 5 EEF in experiment 3 (Fig. 4).

**Figure 4.**
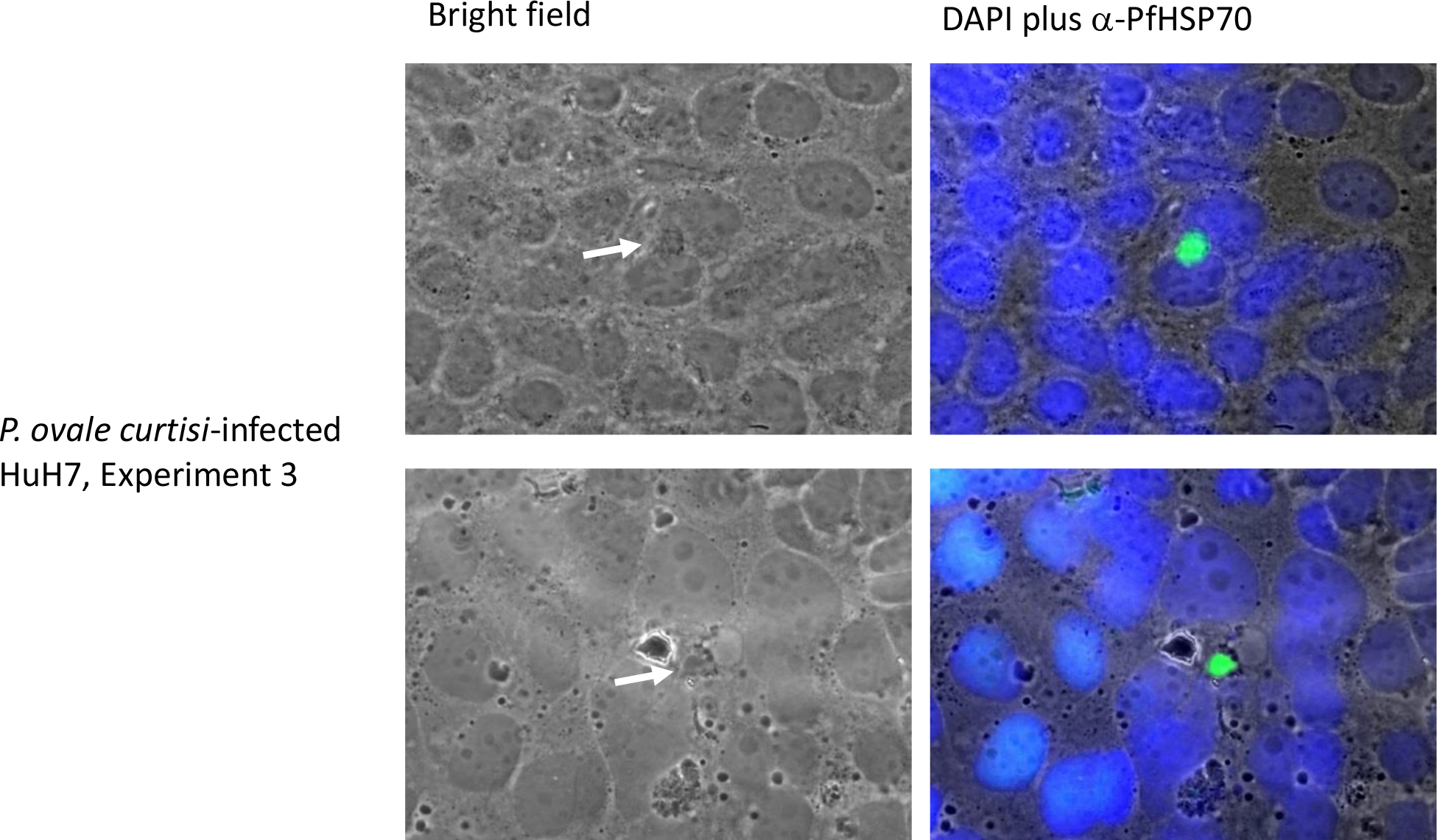
*P. ovale curtisi* extra-erythrocytic forms in human hepatocytes. Sporozoite-exposed HuH7 hepatocytes from experiment 3 were observed by confocal microscopy on day 3. Host and parasite DNA was stained with DAPI (blue). *P. ovale curtisi* EEF (white arrow in bright-field panel) were immunostained with FITC-conjugated monoclonal anti-*P.f* HSP70 antibodies (green).

## DISCUSSION

These results show that routinely collected EDTA-preserved blood samples from patients with imported malaria can be used to successfully infect mosquitoes and produce sporozoites for hepatocyte invasion assays, albeit at low densities. This approach is suitable for further development of an extended protocol for evaluation of drug susceptibility in the hepatic stages of *Plasmodium ovale* spp., and other transmission studies. Importantly, it should now be possible to perform comparative analyses of the hepatic life cycle stages *P. ovale curtisi* and *P. ovale wallikeri*, including assessment of the potential for hypnozoite formation in these *in vitro* systems. This is particularly important as the only demonstrable biological difference, to date, between these two forms of ovale malaria appears to be contrasting delay times to relapse of hypnozoites [7].

There are several reasons to be cautious in making strong conclusions from the results we have presented. Firstly, the lack of material and low efficiency means that repeating and extending the work may be difficult. Usable blood samples from UK cases of infection with *P. ovale* spp. are uncommon; since 2010 between 66 and 130 ovale malaria cases have been reported to Public Health England (PHE) each year (https://www.gov.uk/government/publications/imported-malaria-in-the-uk-statistics), but for only a proportion of these are blood samples of sufficient quality for mosquito feeding sent to the Malaria Reference Laboratory. For the few samples received that were able to be utilised, sporozoite yields from infected mosquitoes were poor, leading to low efficiency in the hepatocyte invasion assays. Further, the blood samples used are not obtained through planned collection but are sporadically received by MRL from a number of UK facilities, posing a number of challenges [7]. They can be a few days old by the time we are able to feed them to mosquitoes, and may not have been kept at optimum conditions for preserving the viability of gametocytes. Further, due to the small sample volume we were not able to wash the erythrocytes to remove EDTA, which is the anticoagulant of choice when collecting venous blood for preparation of thick and thin films for malaria diagnosis [16]. It is known that the presence of EDTA causes some retardation of *P. vivax* development in mosquitoes, but this inhibition is not complete [17]. Reducing EDTA concentration with an equal volume of fresh whole donor blood was used as a simple strategy to minimise any inhibition and to help maintain mosquito feeding rates.

Scaling up will be a challenge with which we will have to deal in the future. The optimum solution would be to conduct active sampling specifically for the purpose of membrane feeding the mosquitoes, using heparin instead of EDTA as anticoagulant [17]. Infected blood should be maintained at 37°C and fed to mosquitoes as soon as possible after collection, to prevent exflagellation and ookinete formation in the tube. Another possibility would be to cryopreserve clinical isolates of *P. ovale* spp. It has been shown that *P. vivax* gametocytes remain stable during the cryopreservation and thawing processes and can be successfully used for sporozoite production, which might be an avenue worth exploring for *P. ovale* spp. as well [18].

Apart from increasing the yields it is also important to enhance the longevity of dissected sporozoites, which would improve the invasion of hepatocytes and development of EEF. Many different media have been used for salivary gland dissection and sporozoite harvesting in the past, and RPMI – as used in this study – is a standard medium for such procedures. However, it was recently shown that using modified Grace’s insect medium significantly increases viability of *P. vivax* and *P. falciparum* sporozoites [19]. Using compatible vectors and parasites can also determine the success of sporogony. All three samples that resulted in infected mosquitoes were of Nigerian origin, matching the parasite to its natural vector (*An. coluzzii* N’gousso strain, originally from Cameroon), which may increase prevalence and intensity of infection [18]. Mosquito survival following the infective feeds was rather low (32.9 – 40.4%), with the highest mortality during the first 5-6 days after the feed. It is still not known exactly what effect malaria parasites have on mosquito survival but it is possible that the parasites incur fitness costs on their vectors [20]. Low survival could also be due to the condition of the original blood sample used as a blood meal.

A number of different *in vitro* platforms for studies of *P. vivax* liver stages have been developed [18], but very little work on *P. ovale* spp. liver development has been published. It has been shown that primary cultured human hepatocytes best support *in vitro* development of *P. ovale* spp. parasites, allowing for sporozoite penetration and further development of EEFs [21]. More recently, humanized mice have been successfully used to observe the development of *P. ovale* liver stages, including hypnozoites [22]. Our experiments show that Huh7 cells, which are derived from human cellular carcinoma cells, also support development of *P. ovale* spp EEFs.

Having access to *P. ovale* spp. patient samples is invaluable and serves as the basis for development of our transmission protocol. Further methodological improvement is now needed to achieve better infection rates and survival of mosquitoes, and therefore more efficient invasion of hepatocytes and development of EEF. These studies provide an important opportunity to study the development, growth, differentiation and drug susceptibility of extra-erythrocytic life cycle stages of *P. ovale curtisi* and *P. ovale wallikeri*.

